# Human sensorimotor beta event characteristics and aperiodic signal are highly heritable

**DOI:** 10.1101/2023.02.10.527950

**Authors:** K. Amande Pauls, Elina Salmela, Olesia Korsun, Jan Kujala, Riitta Salmelin, Hanna Renvall

**Affiliations:** Department of Neurology, Helsinki University Hospital and Department of Clinical Neurosciences (Neurology), University of Helsinki, 00029 Helsinki, Finland; BioMag Laboratory, HUS Medical Imaging Center, Helsinki University Hospital, 00290 Helsinki, Finland; Organismal and Evolutionary Biology Research Programme, Faculty of Biological and Environmental Sciences, University of Helsinki, 00014 Helsinki, Finland; Department of Biology, University of Turku, 20014 Turku, Finland; Department of Neuroscience and Biomedical Engineering, School of Science, Aalto University, 02150 Espoo, Finland; Department of Psychology, University of Jyväskylä, 40014 Jyväskylä, Finland

## Abstract

Individuals’ phenotypes, including the brain’s structure and function, are largely determined by genes and their interplay. The resting brain generates salient rhythmic patterns which can be characterized non-invasively using functional neuroimaging such as magnetoencephalography (MEG). One of these rhythms, the somatomotor (‘rolandic’) beta rhythm, shows intermittent high amplitude ‘events’ which predict behavior across tasks and species. Beta rhythm is altered in neurological disease. The aperiodic (‘1/f’) signal present in electrophysiological recordings is also modulated by some neurological conditions and aging. Both sensorimotor beta and aperiodic signal could thus serve as biomarkers of sensorimotor function. Knowledge about the extent to which these brain functional measures are heritable could shed light on the mechanisms underlying their generation. We investigated the heritability and variability of human spontaneous sensorimotor beta rhythm and aperiodic activity in 210 healthy adult siblings’ spontaneous MEG activity. Both the overall beta spectral power as well as time-resolved beta event amplitude parameters were highly heritable, whereas the heritabilities for peak frequency and measures of event duration remained nonsignificant. Interestingly, the most heritable trait was the aperiodic 1/f signal, with a heritability of 0.94 in the right hemisphere. Human sensorimotor neural activity can thus be dissected into different components with variable heritability. We postulate that differences in heritability in part reflect different underlying signal generating mechanisms. The 1/f signal and beta event amplitude measures may depend more on fixed, anatomical parameters, whereas beta event duration and its modulation reflect dynamic characteristics, guiding their use as potential disease biomarkers.

## Introduction

Individuals’ phenotypes are determined to a significant degree by their genetic blueprint, consisting of a multitude of polymorphic genes and their interplay. Genes regulate our structural and functional properties, ranging from the makeup of cells and their products (Barroso and McCarthy, 2019) to system-level brain macrostructure (Geschwind et al., 2002; Peper et al., 2007). Genetic influences also determine functional brain measures which are constant within, but highly variable between individuals. Functional brain imaging, such as electroencephalography (EEG) and magnetoencephalography (MEG), has been successfully applied to quantify the heritability and even the underlying genetic determinants of several functional brain measures (Van Beijsterveldt et al., 1996; Smit et al., 2006; Koten et al., 2009; Renvall et al., 2012; van Pelt et al., 2012).

The brain generates electrical activity with salient rhythmic, but also arrhythmic patterns at rest. This ‘background’ activity is very prominent, with spontaneous rhythmic oscillations across a range of frequencies, including delta, alpha, beta and gamma bands. The spectral power for several frequency bands is highly heritable (Van Baal et al., 1996; Van Beijsterveldt et al., 1996; Smit et al., 2005, Smit et al. 2006; Salmela et al., 2016).

The most prominent spontaneous rhythms are the parieto-occipital alpha rhythm and the somatomotor (rolandic, or mu) beta rhythm (reviewed by Hari & Salmelin, 1997). The neocortical beta rhythm is observed across several mammalian species, including rodents (Sherman et al., 2016) and non-human primates (Haegens et al., 2011; Feingold et al., 2015; Sherman et al., 2016). It is modulated by many perceptual and cognitive functions, including tactile processing (Pfurtscheller et al., 2001; Haegens et al., 2011), motor function (Salmelin and Hari, 1994; Feingold et al., 2015), action perception (Hari et al., 1998; Babiloni et al., 2002) and attention (Van Ede et al., 2011; Sacchet et al., 2015). Human beta band power has been shown to be heritable (Van Beijsterveldt et al., 1996; Smit et al., 2005), and its variability has been linked to a GABA_A_ receptor locus (Porjesz et al., 2002).

Cortical beta band activity occurs in intermittent high amplitude ‘events’ alternating with lower amplitude (Feingold et al., 2015; Jones, 2016). Healthy sensorimotor cortical brain activity is particularly patterned, with high event amplitudes and long intervals (Seedat et al., 2020). The rate of beta events predicts behavior across tasks and species (Shin et al., 2017). The sensorimotor cortical beta rhythm is altered in neurological conditions affecting motor function, such as genetically determined Unverricht-Lundborg disease (Silén et al., 2000), stroke (Laaksonen et al., 2012) and Parkinson’s disease (Vinding et al., 2020; Pauls et al., 2022). Beta rhythm may thus provide a cortical biomarker of both the disease state and its reactivity to treatment (Laaksonen et al., 2012; Pauls et al., 2022).

Besides rhythmic, or periodic, components, MEG power spectra also contain prominent aperiodic (‘1/f’) components with exponential decay characteristics observed in all electrophysiological data (He, 2014). The rhythmic and aperiodic components are important to disentangle as they are probably generated by different mechanisms. Aperiodic signal is believed to represent excitation-inhibition balance (Gao et al., 2017) and is modulated, *e.g*., by brain maturation (McSweeney et al., 2021; Hill et al., 2022; Tröndle et al., 2022), aging (Voytek et al., 2015; Wilson et al., 2022) and several neurological and psychiatric conditions (Molina et al., 2020; Ostlund et al., 2021; Semenova et al., 2021).

Human cortical peak oscillation frequency decreases with increasing cortical thickness and cortical processing hierarchy (Mahjoory et al., 2020). Many MEG signal characteristics probably reflect individual brain functions more closely than static anatomical structures and could thus detect pathology before brain MRI structural changes emerge. Knowledge about the variability and genetic underpinnings of sensorimotor neural activity could shed light on the underlying generating mechanisms and help interpret changes observed in, *e.g*., patient populations with sensorimotor dysfunction. We investigated the heritability and variability of human cortical sensorimotor beta rhythm and aperiodic activity in healthy adult siblings’ spontaneous MEG, which have not been investigated previously.

## Materials and methods

### Subjects

210 Finnish-speaking siblings from 100 families participated in the study (8 families with three siblings, 1 family with four; 148 females [mean ± SD age 29 ± 10 years], 62 males [30 ± 9 years]; 206 right-handed, three ambidextrous, one left-handed). Monozygotic twins were excluded from the study. None of the participants had a history of neurological or psychiatric disorders. The study was approved by the Hospital District of Helsinki and Uusimaa ethics committee, and all participants gave their written informed consent to participate.

### MEG recordings

Spontaneous cortical activity was recorded in a magnetically shielded room with a 306-channel Vectorview neuromagnetometer (Elekta Oy, Helsinki, Finland) that contains 204 planar gradiometers and 102 magnetometers. Head positioning was measured at the beginning of the measurement. Three minutes of data was collected while participants were resting with their eyes open (REST), as well as while they clenched both hands alternatingly about once per second, self-paced, keeping the eyes open (MOT). The MEG signals were band-pass filtered at 0.03–200□Hz and sampled at 600 Hz.

### MEG signal processing and beta event extraction

For suppressing external artifacts, MEG data were preprocessed using the signal space separation method (SSS, Taulu and Kajola, 2005) implemented in MaxFilter software (MEGIN Oy, Helsinki, Finland). Individual MEG recordings were transferred to one subject’s head space using a signal space separation based head transformation algorithm (Taulu et al., 2004, implemented in MaxFilter). Further signal processing was done using MNE-python version 0.22 (Gramfort et al., 2013). After band-pass filtering the data to 2-48 Hz with a one-pass, zero-phase, non-causal FIR filter (MNE firwin filter design using a Hamming window), power spectral density (PSD) was calculated using Welch’s method (MNE’s psd_welch function) with a non-overlapping Hamming window and 1024-point Fast Fourier Transformation.

The subsequent analysis steps are illustrated in **Figure 1**. The data analysis was performed on the 204 gradiometer signals. First, a channel pair with the highest spectral peak in the beta range (‘the peak channel pair’) was selected from the region of interest (ROI) of 15 gradiometer channel pairs per hemisphere centered over the sensorimotor cortices, and the frequency at the power peak noted (‘peak beta frequency’) (see **Figure 1A**). In order to quantify PSD at each recording site, we computed the vector sum of the two orthogonally oriented planar gradiometers at each sensor location (‘vector PSD’):

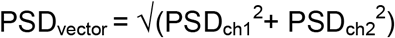

**Figure 1:**
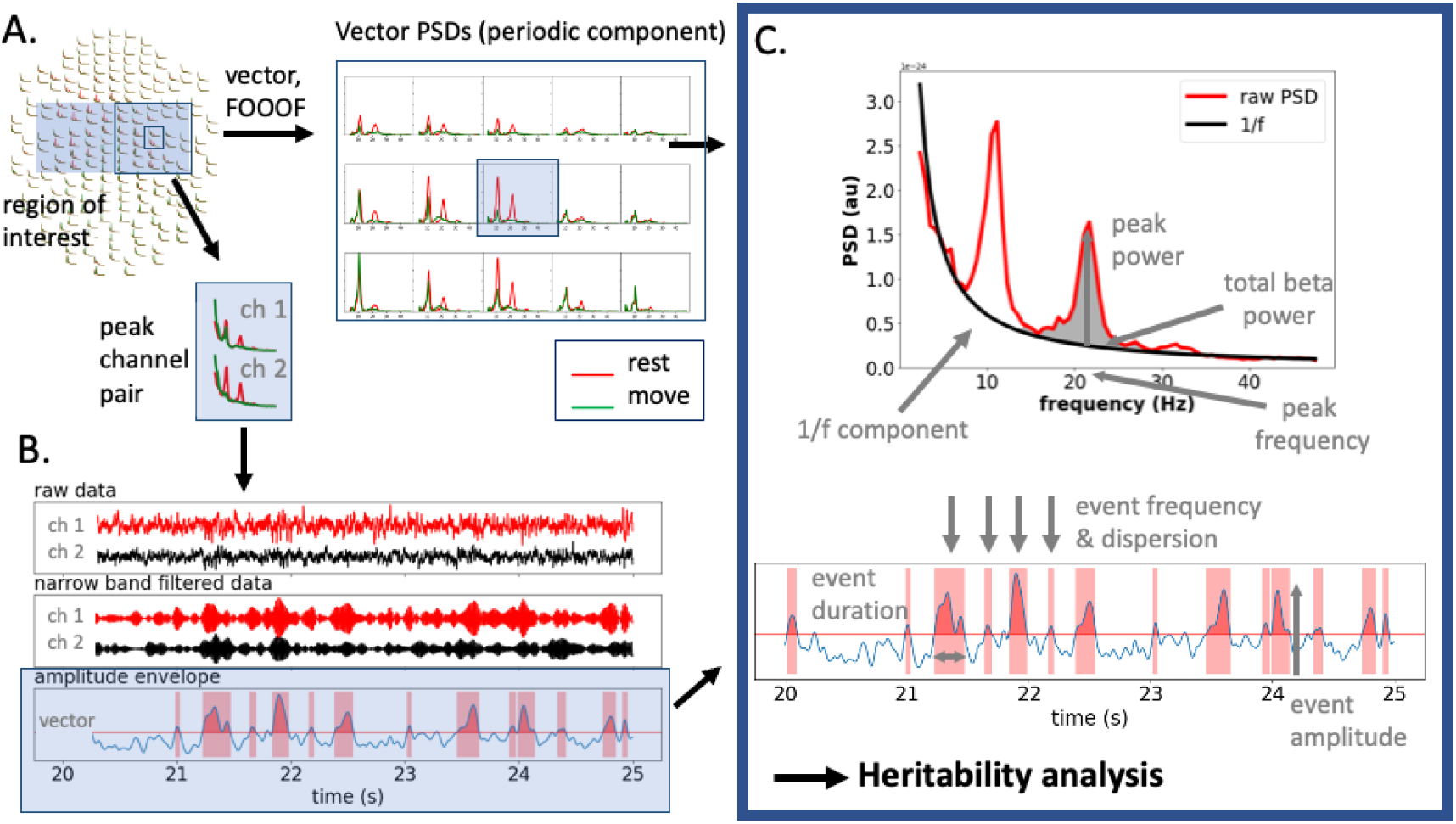
Extraction of sensorimotor beta phenotype characteristics. **A.** Channel selection. A region of interest (ROI) was defined for both hemispheres. The 15 selected gradiometer-channel pairs were combined into 15 vector-sum PSDs (one per channel pair). The periodic spectral component of the vector-sum PSD was obtained using FOOOF. From these, a peak beta frequency and peak channel pair were selected. **B.** Beta event extraction. The peak channel pair and peak frequency selected in A were used to calculate the channel pair’s amplitude envelope. From the raw data, narrow-band filtered data were obtained using wavelet decomposition, and the individual channels’ band-filtered signals were combined to one amplitude envelope using vector sum calculation. **C.** Parameters for heritability analysis. Both PSD characteristics (beta peak power and frequency, total beta power at 14-30 Hz (periodic part), 1/f exponent; upper panel) and time-resolved beta oscillatory characteristics (beta events; lower panel) were used in the heritability analysis.

The resulting 15 vector-sum PSDs per hemisphere were then decomposed into a periodic and non-periodic component using FOOOF (Donoghue et al., 2020). FOOOF models the power spectrum as a combination of two distinct functional processes: an aperiodic component, reflecting 1/f like characteristics (exponential decay with an offset and an exponent), and a variable number of periodic components (putative oscillations), as peaks rising above the aperiodic component. After subtraction of the aperiodic component, the remaining periodic component was plotted for all 15 vector-sum PSDs for both REST and MOT conditions in the frequency range of 14-30 Hz. The resulting plots were visually inspected by two observers (AP and OK) to manually select the beta signal frequency modulated most by MOT compared to REST. Using the manually selected peak frequencies, the periodic components of the 15 vector-sum PSDs were searched automatically to determine the recording channel with the highest peak and its frequency (± 1 Hz) for both hemispheres’ ROIs, and visually inspected again by AP.

The peak beta frequency and corresponding peak power of the chosen vector-sum PSD, the total beta band power (periodic part of PSD area under curve (AUC) from 14-30 Hz, 1/f component subtracted), as well as the non-periodic component information obtained via FOOOF (offset and exponent chi), were further used in the heritability analysis. All parameters included in the heritability analysis are illustrated in **Figure 1C**.

The channel pair and peak beta frequency corresponding to the chosen vector-sum PSD were used for beta burst analysis (see **Figure 1B**). Beta event extraction was carried out similarly to the method described in Pauls et al. (2022): the channel pair’s raw unfiltered time series data were downsampled to 200 Hz, high-pass filtered at 2 Hz and decomposed by convolving the signal with a set of complex Morlet wavelets over the frequency range of 7-47 Hz with 1 Hz resolution and n_cycles=frequency/2. The signal was then averaged within the individual narrow-band beta frequency range, *i.e*., ± 1.5 Hz around the individual peak beta frequency, discarding the other frequencies. The vector sum over the two channels’ beta band time series was calculated as described above, and the resulting signal was rectified to obtain one beta band amplitude envelope for the channel pair. The envelope was smoothed with a 100-ms FWHM kernel and thresholded at the 75th percentile value. Periods exceeding this threshold for 50 ms or longer were defined as beta events. For event amplitude and event duration, the mean, median, robust maximum (defined as mean of the top 5% values) and standard deviation values were calculated. Furthermore, events per second (event rate) and event dispersion were calculated similarly to Pauls et al. (2022). Times between beta events were defined as waiting times. To estimate the variation of waiting times (‘event dispersion’), we calculated the coefficient *C_V_* proposed by Shinomoto et al. (2005), defined as the waiting times’ standard deviation *σ* divided by their mean *μ:*

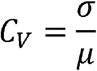

All values were calculated for both hemispheres in all subjects (see also **Figure 1B**).

### Heritability analysis

Phenotype heritabilities were calculated in Merlin version 1.1.2 (Abecasis et al., 2002) using the variance component option. Phenotypes were first transformed by multiplying phenotypes with negative values by −1 and phenotypes with very small values by a suitable factor so that their values would not be too small for Merlin to interpret correctly. Correctness of the input data format was checked by the Pedstats program (Wigginton and Abecasis, 2005). As the analysis assumes the studied phenotypes to be normally distributed while many of them were not, we ran the analyses after first correcting the phenotype values’ distributions using the inverse normal correction internal to Merlin. As both analyses produced highly concordant results, we report here the results based on the noncorrected values.

The probability of the observed heritability values being different from zero was assessed by permuting the family numbers of the study subjects 6000 times and calculating the heritability for each of the permuted datasets. For each phenotype, the number of permutations *k* where the permuted heritability was higher than the heritability observed in the real data was recorded and used to calculate the one-tailed probability of the observed heritability exceeding zero as *k*/6000. This permutation scheme may slightly inflate the permuted heritabilities, as it does not explicitly ensure that the permutation does not reproduce any of the original sibships. This may lead to conservative significance estimates. Likewise, to correct for the multiple tests performed (n = 30), we performed a Bonferroni correction, which may be overly conservative considering that some of the phenotypes were correlated.

## Results

A summary of the beta band phenotypic features (both beta PSD features as well as beta band burst characteristics) is given in **Table 1**. **Figure 2A** shows examples of different beta power spectral phenotypes observed, and **Figure 2B** depicts beta band phenotypes for pairs of siblings. Typical PSD phenotypes were *i*) ones with a narrow peak on either side of 20 Hz, *ii*) a broad band activity typically spanning 15-25 Hz, and *iii*) two distinctive peaks, one typically in the lower beta range (14-20 Hz) and the other in the high beta range (20-30 Hz).

**Table 1.**
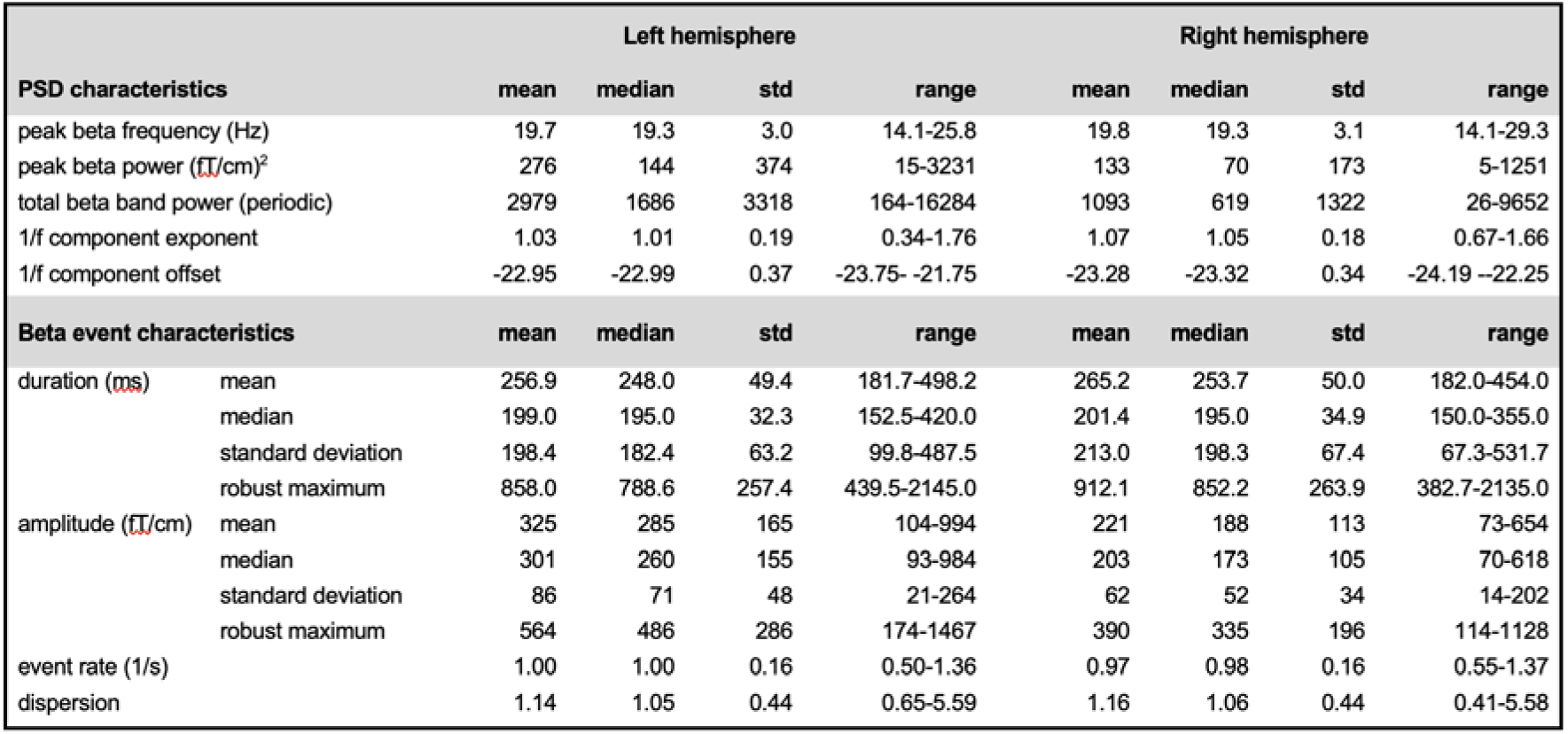
PSD (beta & 1/f) and beta event descriptives. Parameters used in the heritability analysis. Peak frequency – frequency between 14-30 Hz most modulated by hand movement; peak power – PSD amplitude at peak frequency; total beta band power (periodic) – total AUC from 14-30 Hz of the periodic part of the signal (1/f signal component subtracted); 1/f component chi – exponential decay coefficient and offset describing 1/f (aperiodic) signal component. Beta event characteristics: robust maximum – mean of top 5 % values; burst rate – number of bursts/recording time; dispersion – stdev(inter-burst intervals)/mean(inter-burst intervals).

**Figure 2:**
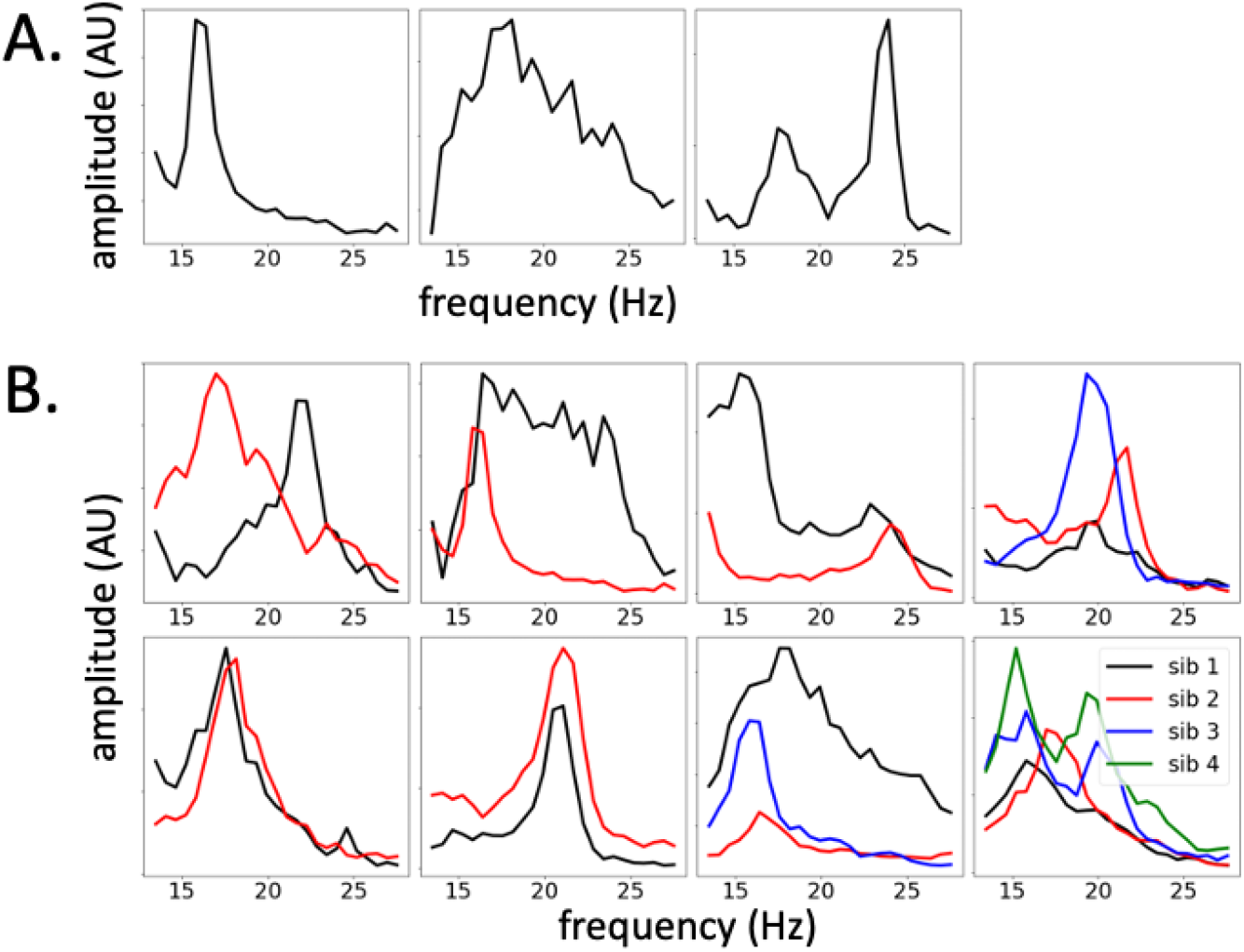
Beta phenotypes. **A. Phenotypic spectrum of beta activity.** Examples of typical beta range PSD patterns: (a) narrow beta peak, (b) broad range, ‘beta brush’ like activity, (c) double peaks of comparable strength, one in the lower, one in the higher beta range. **B. Beta PSD patterns in siblings.** Examples of siblings’ beta PSD patterns (two families with two siblings, one family with three siblings, one family with four siblings).

Heritability results are shown in **Table 2**. Overall, the right-hemispheric parameters were more heritable than the left-hemispheric ones. The 1/f aperiodic exponent was significantly heritable in both hemispheres (right hemisphere h^2^=0.94, left h^2^=0.55). Peak PSD amplitude was significantly heritable on the right (h^2^=0.62), as well as the measures of beta burst amplitudes (heritability h^2^ range 0.65-0.91). Notably, measures reflecting the dynamic range (beta event amplitude maximum and its standard deviation) were most highly heritable.

**Table 2.**
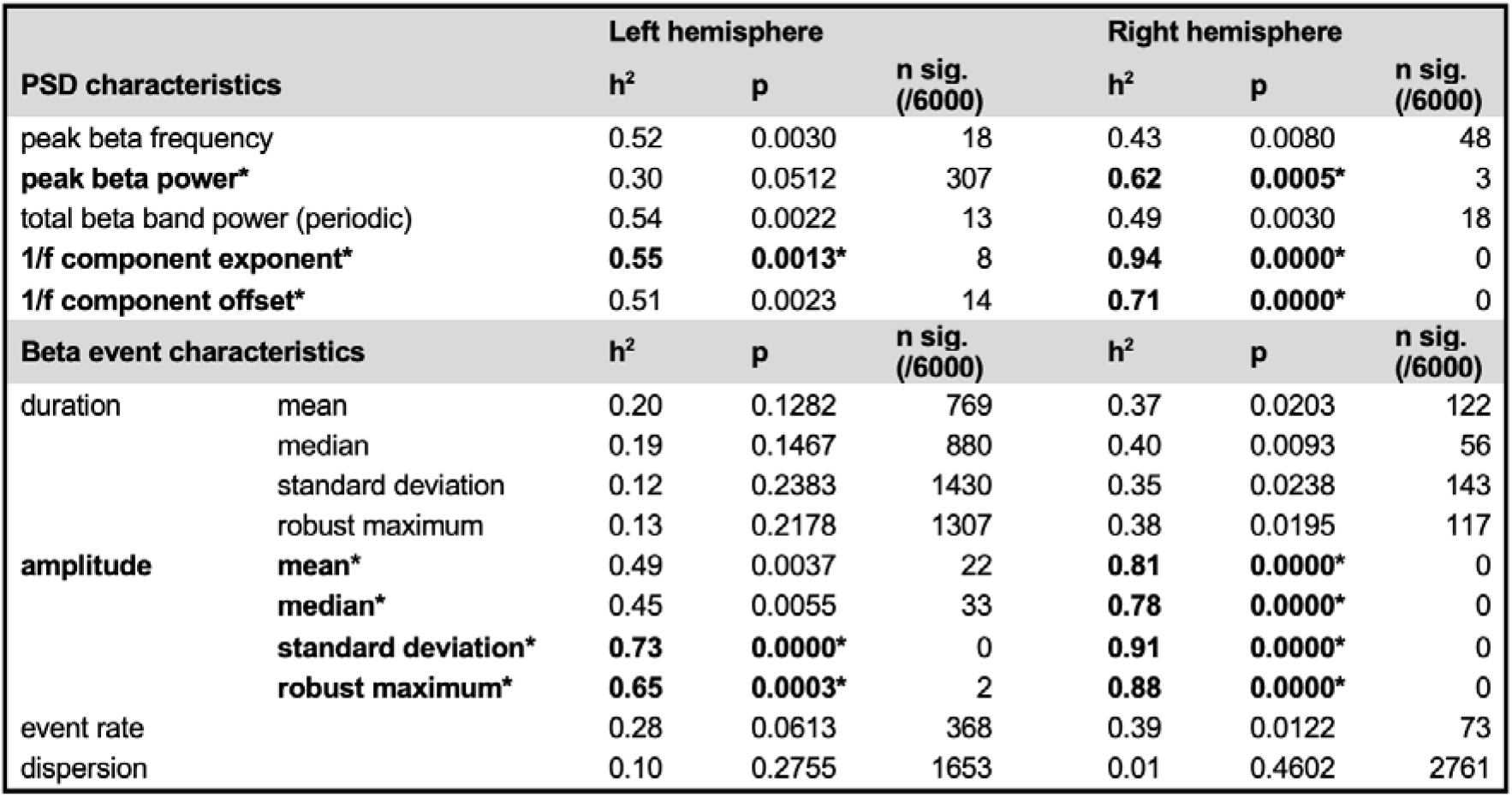
Heritability h^2^ of the oscillatory phenotypes calculated by Merlin. The nominal probability that the heritability differs from zero is calculated from an empirical distribution based on 6000 permutations of the sibship statuses/family IDs of the subjects. The variables and values that are significant after a Bonferroni correction for multiple testing are given in bold.

## Discussion

To our knowledge, this is the first study investigating the heritability of spontaneous sensorimotor beta event dynamics and aperiodic neural activity. Both the overall beta spectral power as well as time-resolved beta event amplitude parameters were highly heritable, whereas the heritabilities for peak frequency and measures of event duration were not significantly different from zero. Interestingly, the most heritable trait was the aperiodic 1/f exponent, with a heritability of 0.94 in the right hemisphere. Overall, the right-hemispheric phenotypic traits were more heritable than the left-hemispheric ones.

### Heritability of MEG/EEG traits including beta oscillatory activity

Heritability of electrophysiological traits has been little investigated to date. In twin studies, EEG alpha, beta, theta and delta range peak frequencies (Van Beijsterveldt et al., 1996), occipital alpha power and peak frequency at rest (Smit et al., 2006), as well as MEG visual task-related gamma peak frequency (van Pelt et al., 2012) have been found to be highly heritable. We have previously demonstrated that auditory evoked fields’ amplitude (Renvall et al., 2012) as well as occipital resting-state alpha oscillatory activity (Salmela et al., 2016) are heritable in siblings, and that MEG power spectral features at rest allow identification of sibling relationship (Leppäaho et al., 2019). These MEG traits were associated with certain genes (Renvall et al., 2012; Salmela et al., 2016; Leppäaho et al., 2019) but it is likely that most functional brain traits are controlled polygenetically. Furthermore, functional connectivity in theta, alpha and beta bands as measured with MEG appears progressively more similar as the strength of genetic relationship increases (Colclough et al., 2017).

### MEG signal generative mechanisms and possible relation to heritability

MEG measures magnetic fields arising from the *temporal* and *spatial summation* of electric currents occurring in the underlying brain tissue (Buzsáki et al., 2012). We postulate that the MEG dynamical measures investigated here reflect different aspects of MEG signal generation. The upper panel of **Figure 3** schematically summarizes factors that contribute to the generation of MEG signals, and the lower panel indicates how those factors may relate to the functional parameters addressed in this study.

**Figure 3.**
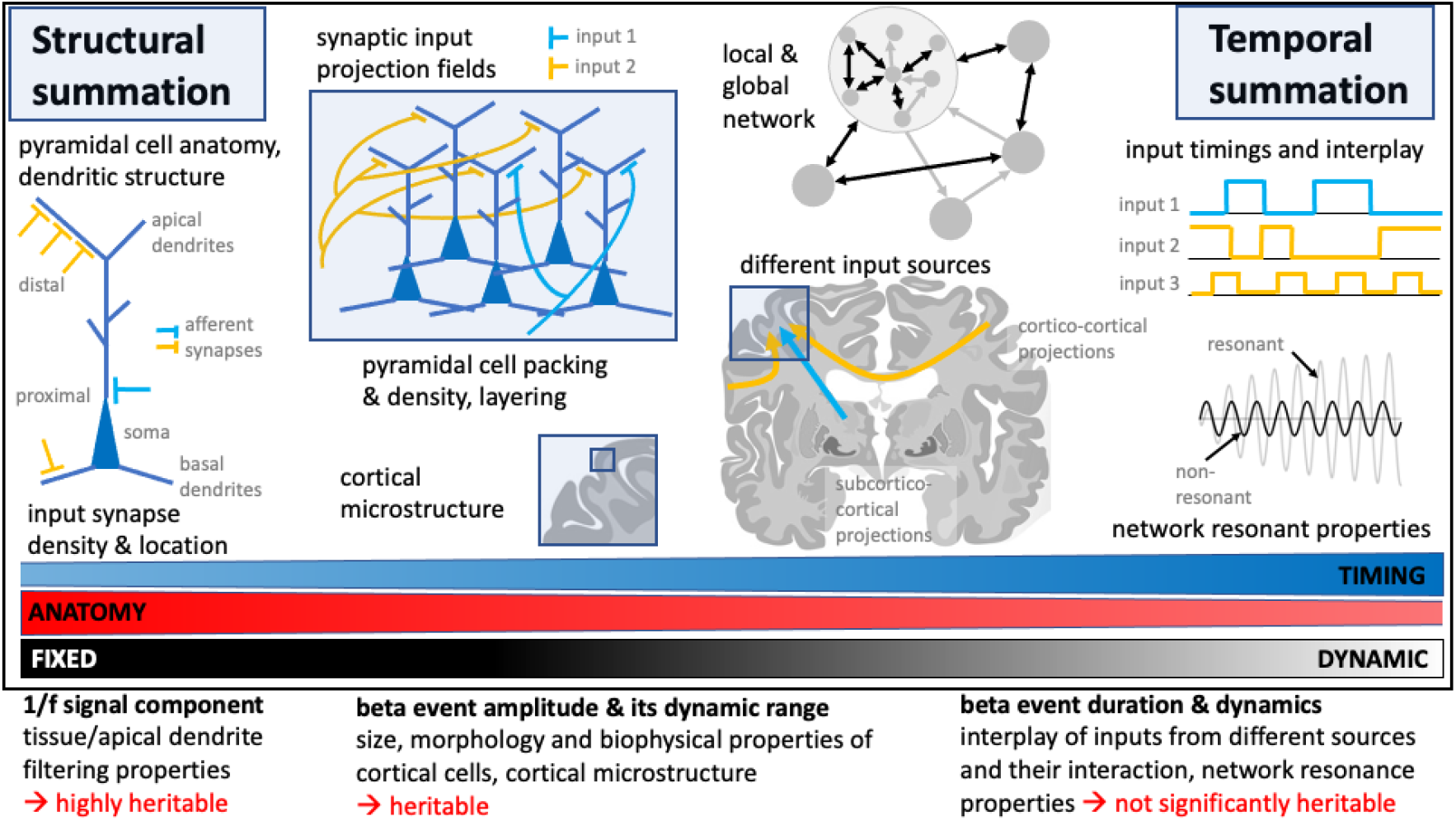
Sources of MEG signals and their putative relationship to the MEG parameters examined in the present study. The schematic figure’s upper panel summarizes the main anatomical and morphological factors, as well as factors determining timing of events, that contribute to the generation of MEG signals. The lower part indicates the putative relationship of those factors to the MEG parameters examined here. We postulate that the 1/f signal and beta event amplitude parameters are more heavily dependent on fixed, anatomical parameters, whereas beta event duration and its modulation are more dynamic characteristics, yet keeping in mind that timing is very much constrained by network anatomy. Brain slice modified from https://commons.wikimedia.org/wiki/File:Human_basal_ganglia_nuclei_as_shown_in_two_coronal_slices_and_with_reference_to_an_illustration_in_the_sagital_plane.svg

### What underlies the heritability of beta event amplitude?

We postulate that the MEG beta event amplitude reflects relatively fixed anatomical factors summarized in **Figure 3** (upper panel, left). Pyramidal cells are neocortex’ most abundant cell type. Synaptic currents and their state-dependent modulation are the main determinants of intra- and extracellular field strength, and their *spatial summation* is governed by pyramidal cell morphology, cortical microstructure and layering, as well as synaptic input density (Buzsáki et al., 2012). Beta event amplitudes are probably crucially dependent on these microstructural properties: While both temporal and spatial superposition determine event amplitude, especially the amplitude’s dynamic range is limited by local cortical microstructure. Interestingly, in the current study, event amplitudes’ dynamic range measures (standard deviation, maximum) were most strongly heritable.

Brain anatomical traits such as cortical thickness (Geschwind et al., 2002; Schmitt et al., 2014) and cortical myelination (Schmitt et al., 2020), have previously been shown to be heritable. By late adolescence, differences in cortical thickness in the sensorimotor regions are largely due to heritable factors, whereas environmental factors play only a weak role (Schmitt et al., 2014). Thus, throughout development, sensorimotor cortical structure appears increasingly governed by the underlying genetics.

Both beta peak amplitude and the power at the beta band (which is determined by the amplitude, number and duration of individual beta events) appeared particularly heritable in the right than left hemisphere. Our result is in agreement with earlier studies that have found cortical morphology/volume to be more genetically controlled in the right than left hemisphere in right-handed individuals (Geschwind et al., 2002); functional studies point in the same direction (Smit et al., 2006).

### Why are event duration parameters not similarly heritable?

In the current cohort, measures of beta event duration were not significantly heritable. *Temporal summation* of neural events, which determines the timing and duration of beta events, arises from the interplay of several brain areas, their connections and relative input timings and strength (**Figure 3**, top panel, right). Important cortical pyramidal cell afferent inputs originate from other adjacent pyramidal cells (intrinsic input) (Lorente de No, 1949), cortico-cortical connections (Kandel et al., 2000) and thalamic connections, including connections from sensory organs, and from other cortical areas (‘higher-order’ thalamic input) (Sherman et al., 2016; Mo and Sherman, 2019). Computational models suggest that sensory induced beta events are generated by synchronous bursts of excitatory synaptic drive to superficial and deep cortical layers, with asymmetry in the respective input strengths (Jones et al., 2009; Sherman et al., 2016; Neymotin et al., 2020): The stronger the superficial input, the more prominent is the beta activity (Sherman et al., 2016). Experimental data are compatible with this model (Sherman et al., 2016; Bonaiuto et al., 2021; Law et al., 2022). Thus, beta event timing and duration appear to depend on the timing and strength of inputs from several different cortical and subcortical input sources,

Network resonance could even play a role in beta event generation: In a dopamine-depleted state, cortical beta events are associated with increased synchrony between EEG/ECoG cortical activity and basal ganglia spiking activity (Cagnan et al., 2019). In animal models of parkinsonism, high cortical beta synchrony can be generated by changing the relative timings between thalamic and cortico-cortical inputs (Reis et al., 2019). Hence, network resonant properties could contribute to temporal summation at least in some disease states, but possibly in a dopamine-dependent fashion also in healthy brains.

Thus, compared to spatial summation, temporal summation relies on more individual factors and their interplay (e.g. network structural and functional properties), making heritability more multifactorial and thus less likely to show heritability in the present analysis. Methodological factors could also contribute to the lack of heritability: signal-to-noise ratio of the recordings affects event duration more than event amplitude measures. Finally, the resting-state beta event duration could be a randomly fluctuating parameter, governed by stochastic events and their timing. These explanations, however, seem less likely given the experimental evidence above.

### Why is the aperiodic signal component heritable?

Aperiodic signal components were the most heritable of the investigated parameters in the present study. The aperiodic signal is closely related to anatomical microstructure: Cortical pyramidal cells and their dendritic morphology and density are believed to be the most important determinants of the mammalian cortical 1/f signal observed with MEG (Lindén et al., 2010; Buzsáki et al., 2012). The 1/f signal is thought to stem from passive dendrite filtering properties (Halnes et al., 2016) but it is also modulated in an activity-dependent way (Pettersen et al., 2014). It has been shown to be affected by processes such as brain maturation (McSweeney et al., 2021; Hill et al., 2022; Tröndle et al., 2022) and aging (Voytek et al., 2015; Wilson et al., 2022) as well as in neurological (Semenova et al., 2021) and psychiatric diseases (Ostlund et al., 2021). Furthermore, 1/f reflects the attentional state (Waschke et al., 2021) and may contribute to integration of signals over longer periods of time (Maniscalco et al., 2018). Thus, the signal’s relative stability over extended periods of time, and its close relationship to cortical microstructure may explain the high heritability.

As for the heritability analysis, the fact that many of the phenotypes were non-normally distributed may have decreased the power of the analysis, as it assumes normal distribution of the phenotypes. Thus, some phenotypes may in fact have higher heritabilities than reported here. Meanwhile, the permutation procedure adopted for testing the significance of the heritability values should correct for any inflation of the heritabilities caused by the non-normality. The analyses were also performed with the internal normality correction functionality of Merlin, and the results were mostly qualitatively similar to (although slightly more significant than) those based on the non-corrected data that were presented here.

## Conclusion

Here, we show that the human sensorimotor beta and aperiodic cortical activity can be dissected into highly heritable and non-heritable components. We postulate that the different heritabilities, in part, reflect different underlying signal generating mechanisms and their weighting in the generation of different signal characteristics. Knowledge of both parameter variability as well as of their heritability are crucial when considering the potential of whole-brain electrophysiology measures, *e.g*., as disease biomarkers.

## Acknowledgements & funding source

We thank all subjects for participating in the study. We acknowledge the following funding sources: AP is funded by a personal grant from Helsinki University, and a grant from the Academy of Finland (grant number 350242). ES has received funding from Jenny and Antti Wihuri Foundation and Ella and Georg Ehrnrooth Foundation. OK is funded by the Instrumentarium Science Foundation. RS has received funding from the Academy of Finland (grant number 315553) and the Sigrid Jusélius Foundation. HR has received funding from the Academy of Finland (grant numbers 127401 and 321460) and the Finnish Cultural Foundation.

